# A Metagenomic Analysis of the Fecal Microbiota of the Critically Endangered Eastern Bongo

**DOI:** 10.1101/2024.06.22.600212

**Authors:** Suzanne M. Kristopeit, Kathryn A. Murphy, Durward L. Bevis, Janelle Brandt, Richard William McLaughlin

**Affiliations:** School of Liberal Arts & Sciences, Gateway Technical College, Kenosha, WI 53144, USA; Great Plains Zoo, Sioux Falls, SD 57104, USA

**Keywords:** Eastern Bongo, fecal material, metagenome, resistome, *Tragelaphus eurycerus isaaci*

## Abstract

The Eastern Bongo (*Tragelaphus eurycerus isaaci*) is a critically endangered mammal. In zoos, this animal has been known to have a sensitive gastrointestinal tract. Using a metagenomic approach the purpose of this study was to determine the microbiota of two adult (n=2) Eastern Bongos living in a zoo in South Dakota, as well as a juvenile (n=6) over a six-week period to see the microbial succession, and to learn if there were pathogenic microorganisms present which are capable of causing gastrointestinal disease. The dominant phylum in seven samples was Bacillota with Pseudomonadota dominant in only one of the juvenile samples. Functional classifications showed Protein Synthesis was the most predominant (11.36–35.71%). Almost equally predominant were Energy and Precursor Metabolites Generation (7.00-13.27%) and Stress Response, Defense and Virulence (8.44-12.90%). Finally, we also determined the resistomes which will contribute to the One Health approach.

## INTRODUCTION

The Eastern bongo, (*Tragelaphus eurycerus isaaci*), is listed in the IUCN Red Data Book as a critically endangered species, with only 70 to 80 mature individuals remaining in the wild (IUCN, 2017). There is actually a greater number of animals living in captivity than in the wild. Unfortunately, in zoos this animal has been known to have a sensitive gastrointestinal tract, commonly resulting in diarrhea (Heng et al., 2021) which can be a clinical sign of enteritis. This condition can be the result of decreased absorption within the intestine (Gunn et al., 2009). In adult ruminants, acute onset hemorrhagic diarrhea can be caused by infectious diseases (Dennison et al., 2002; Songer, 2009).

The gut microbiomes of animals are very important to their health and fitness (Ley et al., 2008; McFall-Ngai et al., 2013). Numerous studies have documented that the gut microbiota plays an important role in various physiological functions, such as protective, structural, and metabolic (Celi et al., 2017). In particular for most ruminants, it has been shown that they are highly reliant on their very diverse gut microbial communities to digest food (Weimer, 2015; Newbold and Ramos-Morales, 2020). Although it has been documented that there is regional diversity along the different parts of the gastrointestinal tract (Suchodolski, 2005; Ritchie et al., 2008; Martinez-Guryn et al., 2019), fecal samples are more practical to collect. In ruminants it has been shown that bacterial communities within the fecal material did not accurately reflect the rumen digesta (Liu et al., 2016; Mohammadzadeh et al., 2014).

However, an accurate characterization of the fecal microbiota was still done as a first step to characterizing the intestinal microbiota.

Exposure to antibiotics can greatly alter the gut microbiota of animals (Tsukayama et al., 2018; Vasco et al., 2023). These chemicals can kill/inhibit the growth of numerous species of microbes which allows for the enrichment of antibiotic resistance genes (ARGs) from the survivors. Ultimately, this allows for the selection of antibiotic resistant bacteria in a particular environment (Uddin et al., 2021). However, even in natural environments which are free of antibiotics produced by humans, bacteria containing ARGs can still be found (Berglund et al., 2015). This could be the result of some species of bacteria and fungi secreting antibiotics into the environment over long periods of time (D’Costa et al., 2011). Unfortunately, today due to the misuse and overuse of antibiotics, resistance has been promoted (Wright, 2010; Allen et al., 2010). As a result, a One Health approach is needed to assess and manage the numerous problems that are caused by resistant bacteria (Robinson et al., 2016).

Despite the status of the bongo as a critically endangered species and reports of enteritis in zoo animals, knowledge of their gastrointestinal tract microbiota is lacking. As a result, the first objective of this study was to accurately determine the fecal microbiota along with their functional classifications and the resistome. The second objective was to document the changes in the intestinal microbiota of the calf as it aged over a six-week period of time using a metagenomic approach.

## MATERIALS AND METHODS

### Collection of fecal samples

Fecal samples were collected from three Eastern Bongos living in a zoo in South Dakota. Beau (male) was born at Zoo Atlanta Dec 02, 2011, and Zahara (female) was born at Omaha’s Henry Doorly Zoo Jul 25, 2014. Bingo (male) was born at the Great Plains Zoo Feb 28, 2022. The fecal samples from Bingo (n=6) were also collected over a six-week period of time (July 27 to August 31, 2022) and samples from Beau and Zahara were collected July 27, 2022. All samples are collected within 24 h from the barn surface into separate sterile containers. The animals live in an outside exhibit or a barn depending on the weather. The barn stall has food and water for the animals. Stalls are cleaned regularly and are rotated among all that are housed in that barn. However, only one animal at a time is present in a stall. The diet of Zahara consisted of produce treats, wild Herbivore and Alfalfa. The diet of Beau and Bingo was wild herbivore plus plus, rhino browser cubes, ADF-16, beet pulp shreds, fruit or vegetable and alfalfa. Zahara or Beau were not on any medications. However, Bingo was on banamine, an anti-inflammatory drug, July 24th, 2022.

Fecal samples were stored frozen, shipped frozen and were stored in a -40°C freezer until used for experiments.

### DNA isolation and metagenomic analysis

The ZymoBIOMICS™ DNA Miniprep Kit (Zymo Research, Irvine, CA, USA) was used for DNA extraction following the manufacturer’s recommendations using the inner portion of the sample to avoid contamination. In brief, the eight separate fecal samples were resuspended in lysis buffer and mechanically lysed using the MP FastPrep-24™ lysis system. After several steps the DNA was washed, eluted, and purified. Finally, a Qubit (Thermo Fischer Scientific, Waltham, MA, USA) was used to determine the concentration.

Illumina sequencing libraries (Illumina, Inc., San Diego, CA, USA) were prepared. This was done using the tagmentation-based and PCR-based Illumina DNA Prep kit and custom IDT 10bp unique dual indices. Sequencing was performed on an Illumina NovaSeq 6000 sequencer in a multiplexed shared-flow-cell runs, producing 2x151bp paired-end reads. Bcl-convert (v4.1.5) was used for demultiplexing, adapter trimming, and quality control.

Metagenomic analysis of the microbial taxa was done using BV-BRC (https://www.bv-brc.org/). Kraken 2 assigns taxonomic labels to metagenomic DNA sequences using the exact alignment of k-mers. The resistome was also determined using BV-BRC. The metagenomic read mapping service which uses KMA (k-mer alignment) to align reads against antibiotic resistance genes present in the Comprehensive Antibiotic Resistance Database (CARD) was utilized. Default settings were used with the exception of *Homo sapiens* reads which were removed for taxa identification. Metabolic profiling was done by merging raw paired reads. Reads which did not merge were also kept. Using Qseq, hits were matched to a custom transcriptome database created using the taxonomic profile as the template for inclusion. The database was derived from genomes included in the BV-BRC database. Data output with gene ID, functional pathways, and subsystems was generated.

## RESULTS AND DISCUSSION

In total, 68, 566, 573 total read pairs were generated. Using BV-BRC 7, 799, 680 classified reads were generated. The Shannon’s diversity index ranged from 2.014 to 4.228 across all eight samples. For Bingo, the juvenile Bongo, the diversity ranged from 3.980 at the beginning of the study to 2.014 at the end of the study.

As shown in Table 1, the percentage of classified sequences ranged from 88.76-98.71, 0.27-11.95 and 0.03-2.47 over all eight samples for the Bacteria, Archaea, and Eukaryota, respectively. As shown in Table 2, the most predominant phyla were the Bacillota (formerly Firmicutes) followed by Bacteroidota (formerly Bacteroidetes) for seven samples. This is a common finding when fecal samples from other ruminants, such as wild blue sheep (Zhu et al., 2020), horses (Costa et al., 2012), Alpine musk deer (Sun et al., 2020), pre-weaned dairy calves (Oikonomou et al., 2013), and dairy cows living in a managed farm from the Patoki region of Pakistan (Khan et al., 2023) were analyzed. However, for cows living in other regions of Pakistan this did not hold true (Khan et al., 2023). One reason the Bacillota and Bacteroidota are so predominant is that members of these phyla contain large numbers of fiber-degrading bacteria (Mizrahi et al., 2021). In particular it has been shown that members of the phylum Bacteroidota can have genes which encode enzymes to hydrolyze complex plant polysaccharides (Grondin et al., 2017) and are capable of producing short chain fatty acids (SCFAs) (Reichardt et al., 2014). From sample Bingo August 31, the most predominant phylum was Pseudomonadota (formerly Proteobacteria) followed by Bacillota.

**Table 1.**
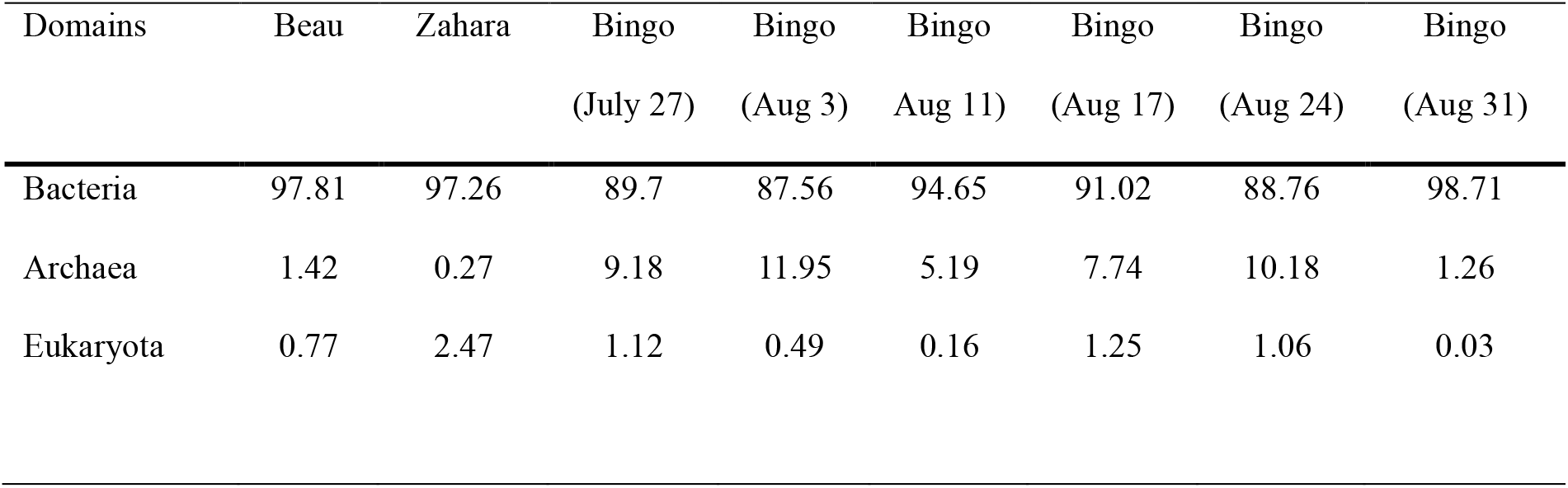
The microbial community composition at the domain level.

| Domains | Beau | Zahara | Bingo | Bingo | Bingo | Bingo | Bingo | Bingo |
| --- | --- | --- | --- | --- | --- | --- | --- | --- |
|  |  |  | (July 27) | (Aug 3) | Aug 11) | (Aug 17) | (Aug 24) | (Aug 31) |
| Bacteria | 97.81 | 97.26 | 89.7 | 87.56 | 94.65 | 91.02 | 88.76 | 98.71 |
| Archaea | 1.42 | 0.27 | 9.18 | 11.95 | 5.19 | 7.74 | 10.18 | 1.26 |
| Eukaryota | 0.77 | 2.47 | 1.12 | 0.49 | 0.16 | 1.25 | 1.06 | 0.03 |

**Table 2.**
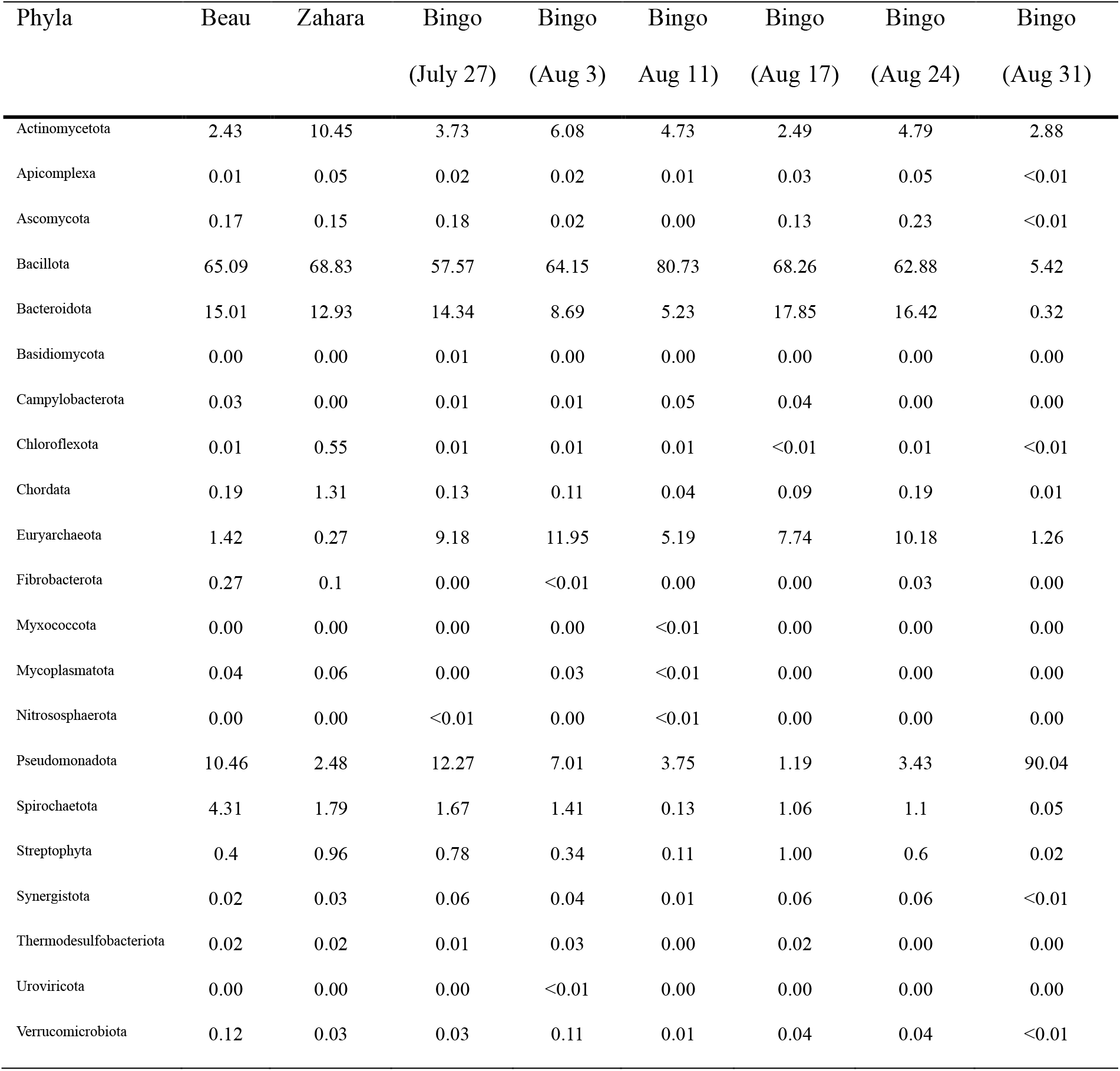
The microbial community composition at the phyla level.

Using rRNA-targeted oligonucleotide probes it was shown that the % SSU rRNA of Archaea found in the colon of steers, goats, and sheep was 0.9, 2.9 and 0.9, respectively (Lin et al., 1997). In our study the Archaea ranged from 11.95 to 0.27% (Table 1). The average Archaea for the two adult Bongos was 1.07%. The average Archaea for Bingo over a six-week period of time was 7.58%. Perhaps juvenile Bongos have a higher percentage of Archaea in their microbiota than adults. All members were part of the phylum Euryarchaeota (0.27 to 11.95%).

### Metabolic profiling

The metabolic profiles for all eight samples were constructed using the subsystem database. Functional classifications showed Protein Synthesis was the most predominant (11.36–35.71%). Almost equally predominant were Energy and Precursor Metabolites Generation (7.00-13.27%) and Stress Response, Defense and Virulence (8.44-12.90%) (Table S2). There were 25 different methane associated genes from 64 different genera of bacteria which were identified (Table S2). Interestingly, in a study using Polish Holstein-Friesian cows in was shown that methane production is only partially controlled by genes (Sypniewski et al., 2021).

### Resistome

The resistome is all of the ARGs present in the bacteria living in a particular environment (Wright, 2007). ARGs move between animals, humans, and different environments (Hu et al., 2017). An improved understanding of antimicrobial resistance in animals as well as other environments has contributed to the One Health approach (Kim and Cha, 2021). In our study, the resistome of the Bongos was examined (Figure 1). In total, there were ten different drug classes, eight which were shared between the two adults. For the juvenile, initially there were nine different drug classes which remained relatively constant with the exception of the August 31^st^ sample which had 35 different drug classes. In total, there were 205 ARGs across the eight samples. The gene sequences are shown in Table S3. The resistome of feed animals, such as livestock and swine, have been extensively studied (Looft et al., 2012; Kim et al., 2023). In studies involving both humans and animals, there is a higher risk of acquiring ARGs among workers involved in the commercial livestock industry. This suggests that the resistome of humans is altered by the resistome of the animals upon exposure thus, highlighting the value of monitoring ARGs in animal husbandry (Van Gompel et al., 2020a; Van Gompel et al., 2020b). However, there are very few studies that examine the resistome of zoo animals which could alter the resistome of zoo workers and the general public upon exposure.

**Fig. 1.**
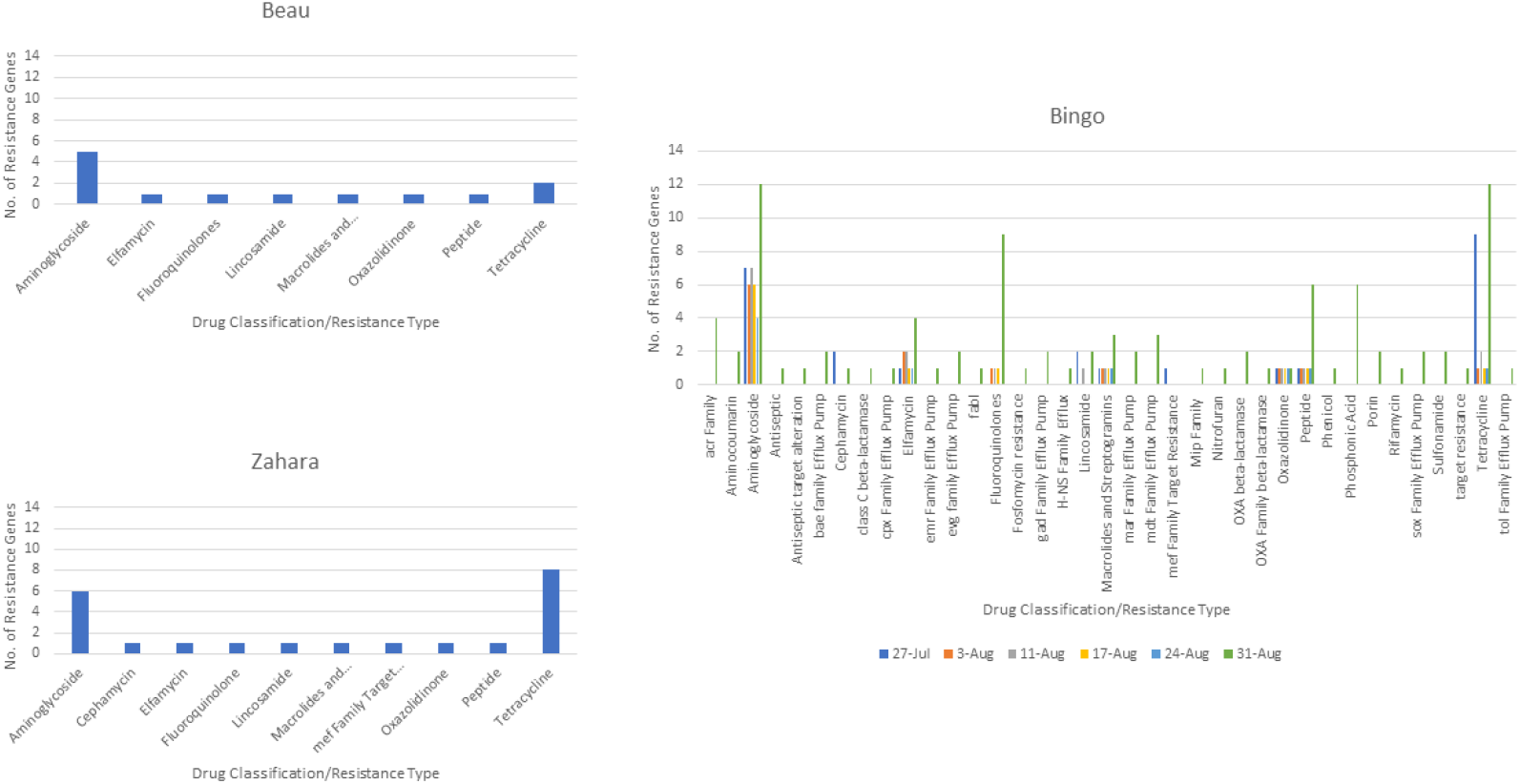
The resistome composition from the fecal material of the Eastern Bongos.

## Supporting information

metagenome analysis

functional classes

antimicrobial resistance

## DATA AVAILABILITY

Sequences have been submitted to the SRA database under accession number PRJNA1082797.

## ACKNOWLEDGMENTS

This research was done as part of a Provost Honors project under the leadership of Matthew Janisin, Executive Vice-President - Academic Affairs. We thank Maxwell Banor, Nicole A.M. Dutton and Donald Zakutansky for their enthusiastic support of this research.

## AUTHOR CONTRIBUTIONS

SMK, KAM and RWM designed this project. JB provided the fecal samples and information about the Eastern Bongos used in this study. SMK, KAM, DLB and RWM were responsible for analyzing data. SMK, KAM, DLB and RWM wrote the manuscript. All authors contributed to discussions, revisions, and approved the final manuscript.

## ETHICS DECLARATIONS

Approval was granted by the Animal Care management team from the Great Plains Zoo for the collection of fecal samples.

## CONSENT FOR PUBLICATION

Not Applicable

